# Bioinformatic analysis of long-lasting transcriptional and translational changes in the basolateral amygdala following acute stress

**DOI:** 10.1101/485888

**Authors:** Stephanie E. Sillivan, Meghan E. Jones, Sarah Jamieson, Gavin Rumbaugh, Courtney A. Miller

## Abstract

Stress profoundly impacts the brain and increases the risk of developing a psychiatric disorder. The brain’s response to stress is mediated by a number of pathways that affect gene expression and protein function throughout the cell. Understanding how stress achieves such dramatic effects on the brain requires an understanding of the brain’s stress response pathways. The majority of studies focused on molecular changes have employed repeated or chronic stress paradigms to assess the long-term consequences of stress and have not taken an integrative genomic and/or proteomic approach. Here, we determined the lasting impact of a single stressful event (restraint) on the broad molecular profile of the basolateral amygdala complex (BLC), a key brain region mediating emotion, memory and stress. Molecular profiling performed thirty days post-restraint consisted of small RNA sequencing, RNA sequencing and quantitative mass spectrometry and identified long-lasting changes in microRNA (miRNA), messenger RNA (mRNA) and proteins. Alignment of the three datasets further delineated the regulation of stress-specific pathways which were validated by qPCR and Western Blot analysis. From this analysis, mir-29a-5p was identified as a putative regulator of stress-induced adaptations in the BLC. Further, a number of predicted mir-29a-5p targets are regulated at the mRNA and protein level. The concerted and long-lasting disruption of multiple molecular pathways in the amygdala by a single stress event is expected to be sufficient to alter behavioral responses to a wide array of future experiences, including exposure to additional stressors.

## Introduction

Stress from external stimuli induces an altered physiological response as the organism compensates to maintain a homeostatic balance [1]. While some stress responses are adaptive to ensure longevity and survival of the organism, maladaptive stress responses can have profound neurological and behavioral effects capable of precipitating the development of psychiatric disorders [2]. Nearly 90% of the population experiences a traumatic stressful event during their lifetime, with higher rates of psychiatric diagnosis positively correlated with the number of traumatic experiences [3]. Indeed, stress is a risk factor for many neuropsychiatric disorders, including schizophrenia, bipolar disorder, posttraumatic stress disorder (PTSD), major depressive disorder, anxiety disorder, and neurodegenerative disorders, that manifest throughout the lifespan of an individual [4; 5; 6]. Stress insults may also prime the brain to develop a more severe phenotype later in life, as individuals that experience early life stress have very high rates of PTSD [5]. Furthermore, patients diagnosed with PTSD as a result of a traumatic event have higher rates of psychotic disorders than non-PTSD patients [7].

Stress induces changes throughout the brain, but a key region involved in stress responses is the basolateral amygdala complex (BLC) [8]. The BLC also mediates learned fear responses and commits fear memories to long-term storage [9]. In the rodent brain, the BLC is one of the most studied subregions of the amygdala in the context of stress because it integrates sensory information related to a potential threat or stressor and conveys this information to the central amygdala, which then activates circuitry to mediate an appropriate motor response to the stressor [10]. Lesions to the amygdala have an anxiolytic effect in both human and rodent studies [11; 12]. Indeed, increased activation of the amygdala has been observed in human studies of individuals with PTSD, panic disorder, social anxiety disorder and other phobia-related anxiety disorders [13]. Human imaging studies have further demonstrated that administration of anxiolytic and antidepressant drugs, which are commonly prescribed to patients with depression, PTSD or anxiety disorders, reduce activation of the amygdala during emotional processing tasks [14; 15], supporting the theory that an overactive amygdala can elicit an anxious phenotype and reducing amygdala activity is therapeutic for stressed individuals. Understanding the lasting impact of stressful experiences on neurobiological function of the BLC, therefore, represents a key step towards designing intervening therapies to combat stress-induced maladaptive responses.

Many different stress paradigms have been employed to study stress-responsive molecular signatures and have demonstrated that stress can elicit both immediate and lasting changes in brain cellular pathways [16]. Because one microRNA (miRNA) can regulate many targets simultaneously, studying miRNAs in the context of complex behavioral phenotypes may yield unique insight into the concerted pathways required to maintain a long-lasting response to a stress event. miRNAs are small non-coding RNAs approximately 20-24 nucleotides in length. They contain a 6-8 nucleotide seed region that allows them to bind to a target region of messenger RNA (mRNA) with sequence complementarity [17]. Once bound, the miRNA triggers deadenylation of the poly-A tail of mRNA, blocking the translation of the mRNA into protein [18]. The short seed region of the miRNA allows for redundancy and binding of one miRNA to many putative target mRNA sequences that can be predicted with web-based algorithms [19; 20; 21]. The miRNA signature that results immediately after stress has been primarily studied in chronic stress paradigms [22], with some focus on immediate effects of acute stress. However, the potential for long-lasting changes in miRNA expression after acute stress is unknown.

Studies to date have shown that varying the strength and duration of a stressor differentially influences the development of maladaptive stress responses [23]. Brief stressors that represent a single traumatic event may have different consequences on brain neurochemistry than chronic stressors and more in-depth analysis of the long-lasting effects of acute stress would provide additional targets for therapeutic treatment of stress disorders. Therefore, it is essential to understand the miRNA pathways that are regulated by acute stressors, in addition to the commonly studied chronic stress paradigms [24]. For example, acute restraint stress of one 2hr session, but not chronic restraint of 2hr sessions for 5 days, immediately induces expression of let-7a, miR-9 and miR 26-a/b in the frontal cortex [25]. Interestingly, expression of mir-34 is elevated in the central amygdala immediately after acute restraint stress and the overexpression of this miRNA in a chronic stress paradigm has anxiolytic effects [26], suggesting that studying acute stress pathways may give insight into stress pathways that can be targeted for alleviation of more severe stress phenotypes as well.

Although chronic stress paradigms are more commonly employed, additional value can be obtained from understanding the extent and duration of impact from acute stress. At the targeted level, a vast number of studies have been done to identify changes in mRNA or protein expression in the BLC that result from stress, but it is highly unlikely that a single protein or gene is responsible for the multitude of phenotypes and neuropsychiatric disorders that arise after traumatic stress exposure. To achieve a systems level network view of stress-responsive pathways, RNA sequencing, RNA microarrays and quantitative proteomics have been performed independently on amygdala tissue after various forms of chronic stress [27; 28; 29]. Such studies have highlighted the involvement of synaptic glutamate signaling in mediating plasticity of the amygdala after 8 weeks of chronic stress and identified networks of genes that are regulated at various timepoints after chronic social defeat, two paradigms that induce symptoms of major depressive disorder in rodents [28]. Similarly, we previously employed a stress-enhanced fear learning paradigm that combines a single acute restraint stress session with Pavlovian fear conditioning and used RNA-sequencing to identify differential expression of networks of genes involved in learning and plasticity in the BLC 30 days after the initial stress event [30]. Non-neuronal networks may also mediate stress responses, as two studies have identified astrocyte marker regulation and oligodendrocyte expression differences in stress paradigms [29; 31].

While these findings demonstrate that stress induces significant changes in multiple molecular pathways in the amygdala, few studies of stress neurobiology have integrated miRNA datasets with RNA-sequencing [32; 33] and none have aligned small RNA sequencing of miRNAs with a corresponding proteomics profile. The latter would be expected to yield highly relevant information, as miRNAs principally repress protein translation. Moreover, the existing datasets in the literature have focused almost exclusively on the effects of chronic stress [22], while the few that have employed acute stress focused on immediate effects and give no insight into the long-term consequences of a brief stress exposure [26; 34]. To address these limitations, we used sequencing and mass spectrometry technologies to identify long-lasting neuroadaptations in the BLC that are regulated by a single stressful event. By combining multiple methods that measure global expression of RNA and protein within the BLC, we have used an unbiased approach to systematically identify miRNA-dependent and -independent pathways at both the RNA and protein level that are changed 30 days after a single stress event. We highlight mir-29a-5p as a key stress-regulated miRNA that has many putative target pathways regulated in the opposite direction, suggesting that mir-29a-5p may modulate long-lasting neuroadaptations in the BLC after stressful experiences.

## Results

### Acute stress regulates expression of miRNAs in the BLC

To determine the long-lasting effects of a single stress event, we measured the molecular consequences in the BLC of restraint after 30 days (Fig 1A).

**Fig 1.**
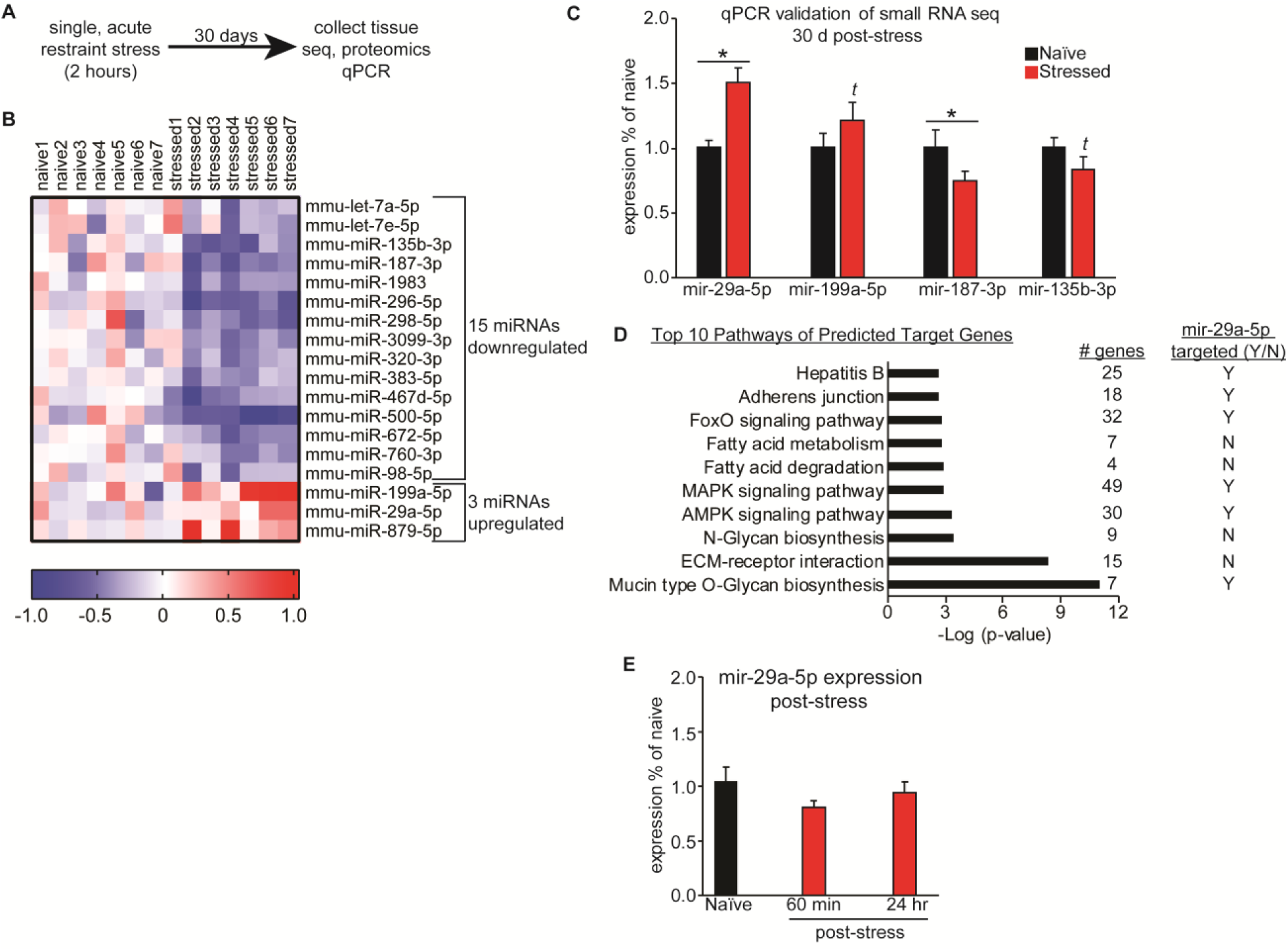
Acute restraint stress induces long-lasting changes in BLC miRNA expression. (A) Overview of the experimental timecourse. (B) Heatmap of the 18 miRNAs differentially expressed in the BLC of mice that went through acute restraint stress 30 days earlier. Red colors indicate upregulation and blue indicate downregulation of a given miRNA. (C) qPCR validation of the smRNA-seq was performed for 3 miRNAs. *t* indicates a trend in the same direction as the smRNA-seq. * indicates p<0.05. Error +/-s.e.m. (D) The top 10 significant pathways of predicted target genes for the list of stress induced miRNAs obtained using DIANA’s miRPATH software. To the right of each pathway are listed the number of target genes that belong to the pathway and a yes (Y) or no (N) to indicate if the pathway is targeted by mir-29a-5p. (E) qPCR experiments were run to examine mir-29a-5p at 60 minutes and 24 hours post-stress, no significant differences were observed between groups.

Naïve mice were handled for 3 days to reduce their baseline anxiety and then restrained for 2 hours during an acute stress session, which we have previously reported elevates corticosterone release [30]. Animals were returned to their home cage immediately after the stress event, where they remained for 30 days, until tissue collection. High quality RNA (RIN>7.0) and protein were extracted from the BLC from multiple cohorts of animals and used for global miRNAome, proteome and transcriptome profiling. To begin, small RNA sequencing (smRNA-seq) was performed on BLC tissue isolated from previously stressed animals and handled controls. To reduce false positives, we performed sequencing in two biological replicate cohorts (N=3-4/group/cohort) and identified a miRNA profile that was significantly changed in both sequencing runs with a p-value of less than 0.05. 18 miRNAs were differentially expressed compared to naïve mice, with 3 upregulated and 15 downregulated (Fig 1B and Tables SE.1-2). Four miRNAs were selected for qPCR validation in a set of technical replicates that included sequenced animals as well as an additional cohort of animals that were not sequenced as a set of biological replicates (naïve N=16-19; stressed N=11-12) (Fig 1C). We prioritized candidate miRNAs based on their degree of variance between replicates. Upregulation of mir-29a-5p and downregulation of mir-187-3p in stressed animals indicated by sequencing was confirmed by qPCR (two-tailed one sample t-tests, mir-29a-5p: t_(11)_ = 4.207, p = 0.0015; mir-187-3p: t_(11)_ = 3.751, p = 0.0032) and the other two miRNAs, mir-199a-5p and mir-135b-3p, showed a statistical trend in the expected direction based on the sequencing results (two-tailed one sample t-tests-mir-199a-5p: t_(11)_ = 1.457, p = 0.1732; mir-135b-3p: t_(10)_ = 1.623, p = 0.1357). We performed a pathway analysis with DIANA miRPATH’s microCts function to identify KEGG pathways that include target genes of the 18 stress-specific differentially expressed miRNAs. This software analysis first identifies all target genes of a miRNA list and then examines the target gene list to determine if there is significant enrichment for genes in a known KEGG pathway. We identified the top 10 most significantly enriched pathways for the 18 stress regulated pathways and 50% of them included known targets of mir-29a-5p (Fig 1D). Significantly enriched pathways of the regulated miRNA list included several pathways with signaling cascades and transcription factors known to respond to physical or oxidative stressors and to mediate stress responses, such as mitogen-activated protein kinase (MAPK), adenosine-monophosphate-activated protein kinase (AMPK) and forkhead box protein (FoxO) signaling pathways. Though mir-29a-5p has not yet been studied in the brain, the persistent elevation of this miRNA suggests it may participate in amygdala-mediated stress responses and putative regulation of stress-mediated signaling molecules. We subsequently measured BLC mir-29a-5p levels at two earlier time points following the single restraint session to determine if the rise in this miRNA occurs as a rapid or delayed response to stress (Fig 1E). BLC mir-29a-5p is not differentially expressed 60 minutes or 24 hours following acute stress (one way ANOVA, F_(2,21)_ = 1.709, p = 0.2054), providing evidence that the increase emerges at some time between 24 hours and 30 days post-stress and suggesting mir-29a-5p signaling is likely to play an important role in the incubation of stress-induced changes.

### BLC protein expression is changed one month after acute stress

In a separate cohort of animals, we performed quantitative mass spectrometry to identify long-term changes in the BLC proteome after acute restraint stress. We employed a tandem mass tag (TMT) method that allowed for relative quantitation between pooled sample groups of either naïve or stressed animals (n=1/group, representing 8 animals’ samples pooled in each group) and reliably detected expression of ∼ 2,400 proteins (Fig 2A and Table SE.3).

**Fig 2.**
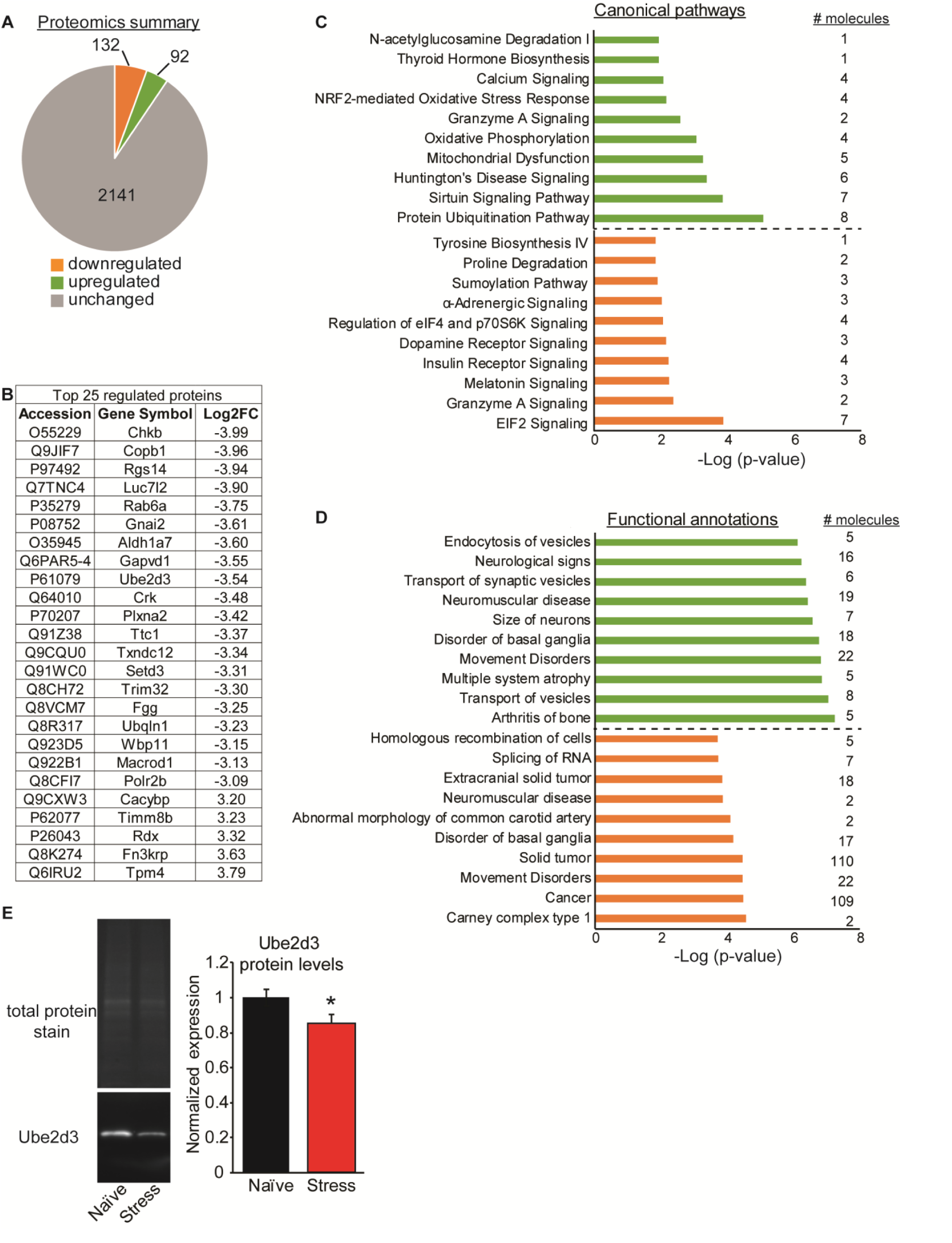
Acute restraint stress induces long last proteomic changes in the BLC. (A) A summary of the global proteomics profile of the BLC in mice that underwent a single acute restraint session 30 days prior. (B) The top 25 most differentially regulated proteins in the BLC of stressed mice and their log2 fold change (Log2FC) values. (C+D) Ingenuity Pathway Analysis of the global proteomics profile identified the top 10 most significant canonical pathways (C) and functional annotations (D) of the gene list. Upregulated pathways are shown in green and downregulated in orange, with the number of molecules belonging to each pathway indicated to the right. (E) Representative images of western blot validation of Ube2d3 protein. On the left are the total protein stain for normalization and the image of Ube2d3. On the right is the quantification of the western blot data.

9.5% of all proteins detected in the BLC were differentially expressed by more than 1.5 log2 fold change between naïve and stressed animals, with 132 downregulated and 92 upregulated. The 25 proteins showing the greatest degree of stress-induced change included proteins known to regulate intracellular stress responses, such as ras-related protein Rab-6A (Rab6a), calcyclin binding protein (Cacybp), plexin A2 (Plxna2), regulator of G protein 14 (Rgs14) and coatomer protein complex subunit beta 1 (Copb1) (Fig 2B). Copb1 is associated with non-clathrin coated vesicles and has been found to complex with mRNA for the kappa opioid receptor (KOR), which plays an integral role in stress responses [35; 36]. To interrogate the stress-responsive proteome, we performed analysis with Ingenuity Pathway Analysis (IPA) software and report the top 10 most significantly upregulated (green) and downregulated (orange) canonical pathways (Fig 2C), as well as functional annotations (Fig 2D). The majority of the pathways identified were unique for the upregulated group relative to the downregulated, with a few exceptions. Both groups of regulated proteins had canonical pathways involved in Granzyme A signaling and functional annotations involved in Disorders of the Basal Ganglia or Movement disorders. This overlap suggests shared functional effects in these major pathways. However, all other pathways were divergent for the up-versus down-regulated protein lists. Because the proteomics represents pooled samples with one pooled sample per group, we selected two proteins for validation, ubiquitin-conjugating enzyme E2D 3 (Ube2d3) and choroideremia-rab escort protein 1 (Chm) in individual replicate samples. These proteins were selected because they exhibited a high fold change value between treatment groups and had published, commercially available antibodies. We measured expression of these two proteins in all samples that were pooled for proteomics, as well as an additional 23 samples, some of which were also used for small-RNA sequencing. The antibody for Chm failed for technical reasons, as no single specific band could be identified by Western blot, but the antibody for Ube2d3 was specific and confirmed downregulation of the Ube2d3 protein in the stress alone group compared to naïve (two-tailed t-test: t_(35)_ = 2.10, p = 0.043, naïve N=21, stressed N=16) (Fig 2E).

### Acute stress has long-lasting consequences on the BLC transcriptome

Finally, we performed RNA-seq on BLC tissue samples from animals that had gone through acute restraint stress 30 days earlier and were also sequenced for small RNAs (naïve N=2, stressed N=4). This traditional method of RNA-seq depletes samples of ribosomal RNA and primarily detects differential expression of mRNA, as opposed to the miRNA changes reported in Figure 1. Due to the size selection process during sequencing library preparation, expression of mature small miRNAs cannot be detected with RNA-seq. We detected expression of ∼13,500 genes in the BLC and 4.7% were significantly changed by acute stress with an adjusted p-value of less than 0.05, including 199 downregulated and 431 upregulated (Fig 3A-B and Table SE.4).

**Fig 3.**
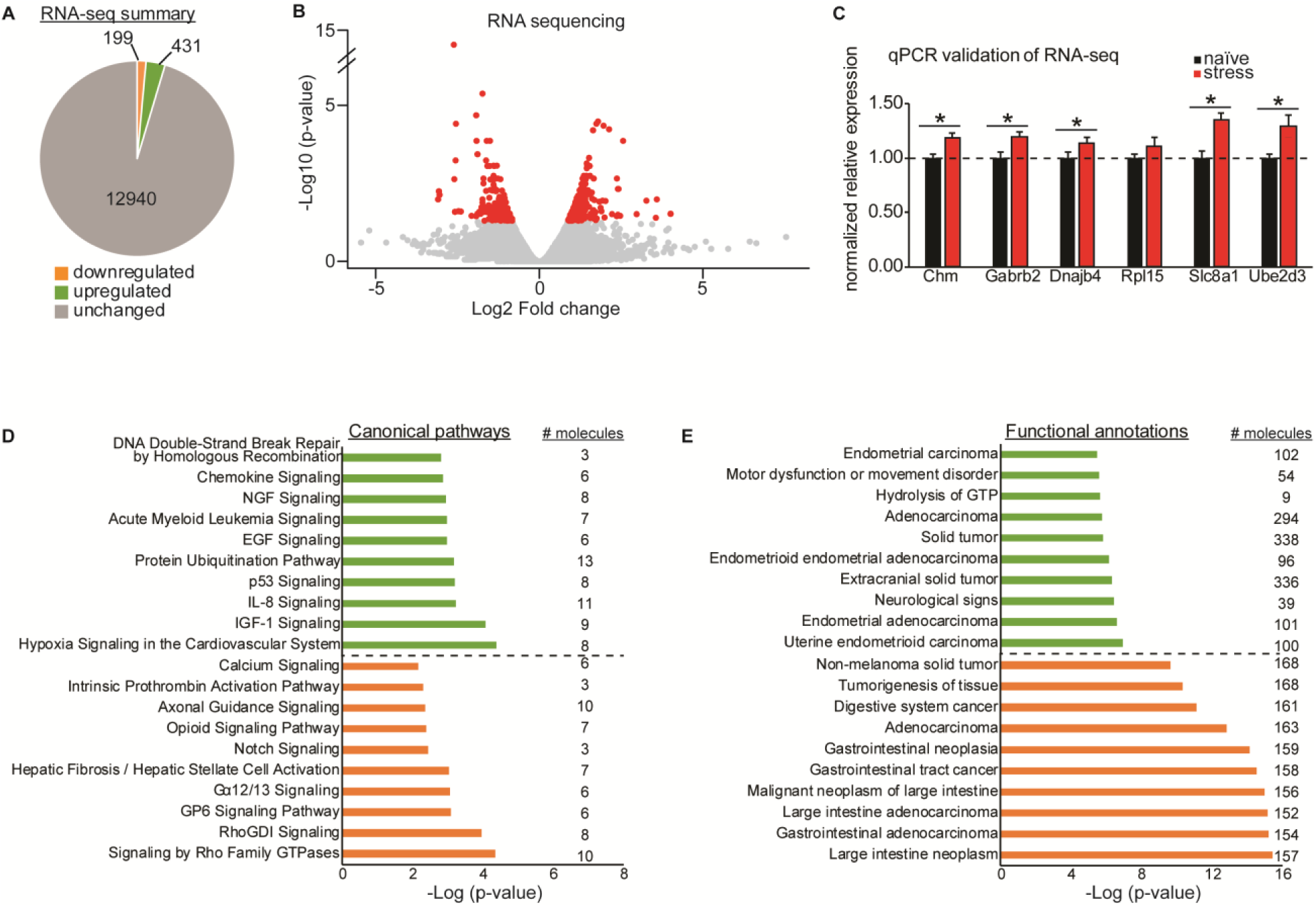
Acute restraint stress induces long lasting RNA changes in the BLC. (A) A summary of the global transcriptome profile of the BLC in mice that underwent a single acute restraint session 30 days prior. (B) A volcano plot of the RNA-seq data with significantly changed genes colored in red and unchanged genes colored in gray. (C) qPCR validation of a subset of genes significantly upregulated in the RNA-seq. *p<0.05. Error +/-s.e.m. (D+E) Ingenuity Pathway Analysis of the global transcriptional profile identified the top 10 most significant canonical pathways (D) and functional annotations (E) of the gene list. Upregulated pathways are shown in green and downregulated in orange, with the number of molecules belonging to each pathway indicated to the right.

We selected 6 genes for qPCR validation that were significantly changed at the RNA level and showed a high degree of fold change at the protein level (>1.5 log 2 fold). We validated these gene changes in technical replicates that included an additional 19 that did not undergo mRNA sequencing (naïve N=12-13; stress N=12) and all displayed an expression pattern consistent with the sequencing results. Five genes had significantly elevated expression in stressed animals relative to naïve (two-tailed one sample t-tests-Chm: t_(11)_ = 5.05, p = 0.0004; gamma-aminobutyric acid type A receptor beta2 subunit (Gabrb2): t_(11)_ = 5.02, p = 0.0004; DnaJ heat shock protein family (Hsp40) member B4 (Dnajb4): t_(11)_ = 2.546, p = 0.027; solute carrier family 8 member A1 (Slc8a1): t_(11)_ = 6.034, p <0.0001; Ube2d3: t_(11)_ = 2.97, p = 0.013) (Fig 3C). IPA analysis of up-or down-regulated genes separately once again revealed divergent canonical pathways for the gene lists (Fig 3D). However, the functional annotations identified by IPA for the differentially expressed RNAs were similar for both up-and -downregulated genes because they both involved many cell signaling pathways traditionally implicated in cancer pathology (Fig 3E). This indicates that genes commonly studied in cancer pathways, including many implicated in cell growth, may be relevant for study in the brain for their role in stress-mediated adaptations.

### Acute stress differentially regulates both miRNA-dependent and independent pathways

We next determined correspondence between mRNA and protein datasets, as often changes in the transcriptome are not ultimately reflected in the proteome. Of the 630 differentially expressed genes at the RNA level, only 138 were technically detected by mass spec in the proteomics dataset. Of the 138 transcripts that could be assessed for corresponding changes at the protein level, 16 were regulated by acute stress at the RNA and protein level (Fig 4A-B).

**Fig 4.**
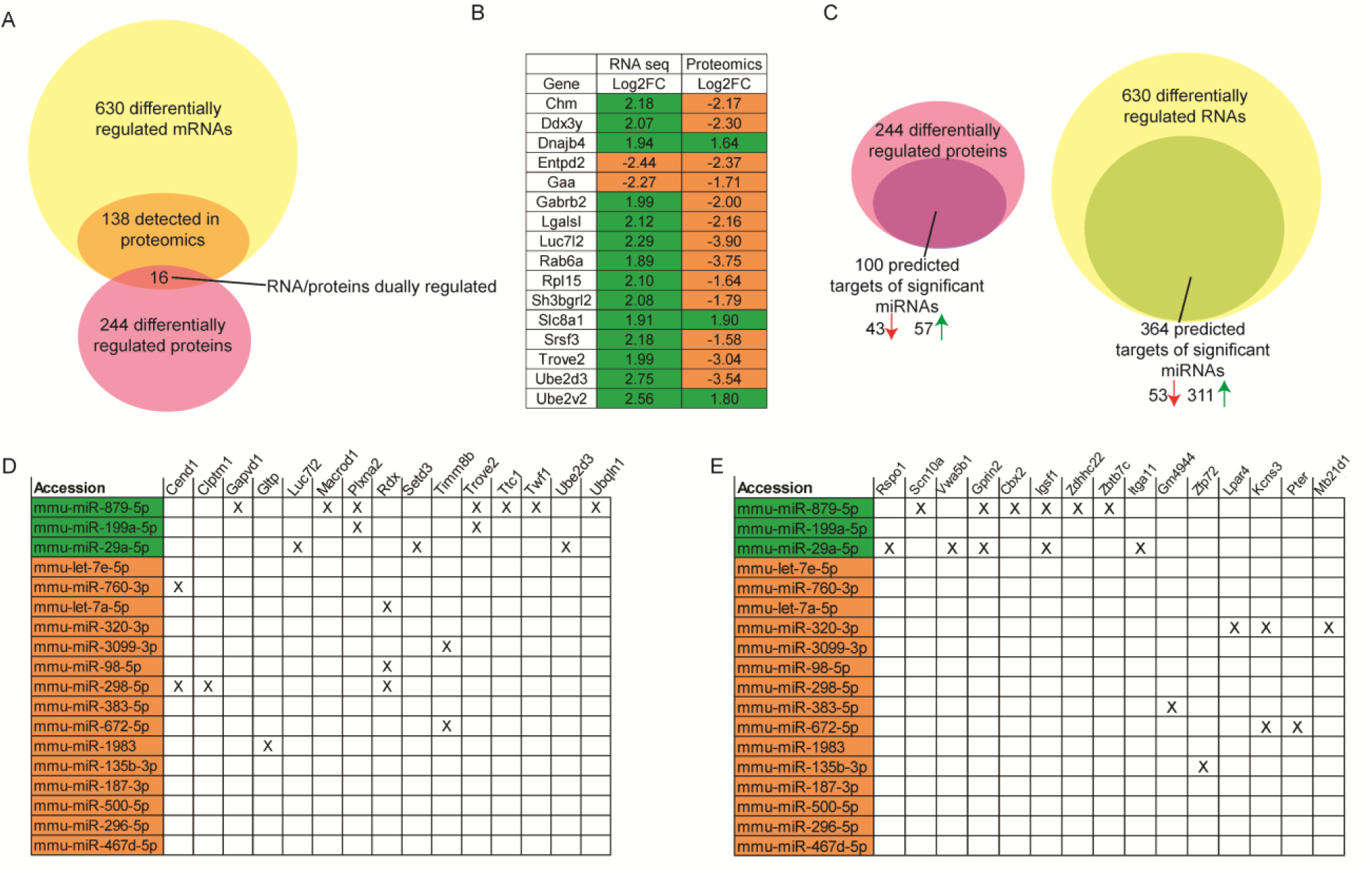
Integration of RNA-seq, smRNA seq and proteomics datasets. (A) Venn diagram displays the overlap of differentially expressed genes at the RNA and protein levels. 16 proteins were regulated at both levels and their log2 fold change (Log2FC) values for each measure are depicted in (B). Upregulated changed are highlighted in green and downregulated changes are highlighted in orange. (C) Integration of the datasets identified putative miRNA targets that are regulated in the opposite direction from significantly changed miRNAs in both the proteomics and RNA-seq datasets. (D-E) Matrices depict the differentially expressed miRNAs that also target the top 15 most regulated genes at the RNA (D) or protein level (E) in an opposing fashion. Upregulated miRNAs are highlighted in green and downregulated in orange.

We next integrated the three datasets by looking at convergence amongst the significantly changed miRNA list and putative targets changed by either proteomics or RNA-seq (Fig 4C). For each of the 18 stress-regulated miRNAs, we made a list of putative target genes from the three major target site prediction websites: TargetScan, mirDB and DIANA. We identified all proteins or mRNAs that were predicted targets of the differentially expressed miRNA list and were regulated in the opposite direction compared to the miRNAs. Interestingly, 42 of mir-29a-5p’s predicted targets were downregulated in the RNA-seq or proteomics analyses (Tables SE.1 and 2). In comparing the miRNA data with the transcriptome and proteome datasets, we identified 100 proteins and 364 mRNAs that were predicted targets of regulated miRNAs (Fig 4C). The protein targets were split with nearly half downregulated and the other half upregulated, despite only 3 miRNAs being upregulated and 15 miRNAs downregulated. For the mRNA list, 53 mRNAs were downregulated but 311 were upregulated. Finally, we integrated the datasets by creating a matrix that displays the top 15 mRNAs or proteins that were the most highly regulated in the RNA-sequencing or mass spectrometry experiments and the significantly regulated miRNAs that are predicted to target them (Fig 4D-E). Interestingly, 42 of mir-29a-5p’s predicted targets were downregulated in the RNA-seq or proteomics analyses (Fig 5).

**Fig 5.**
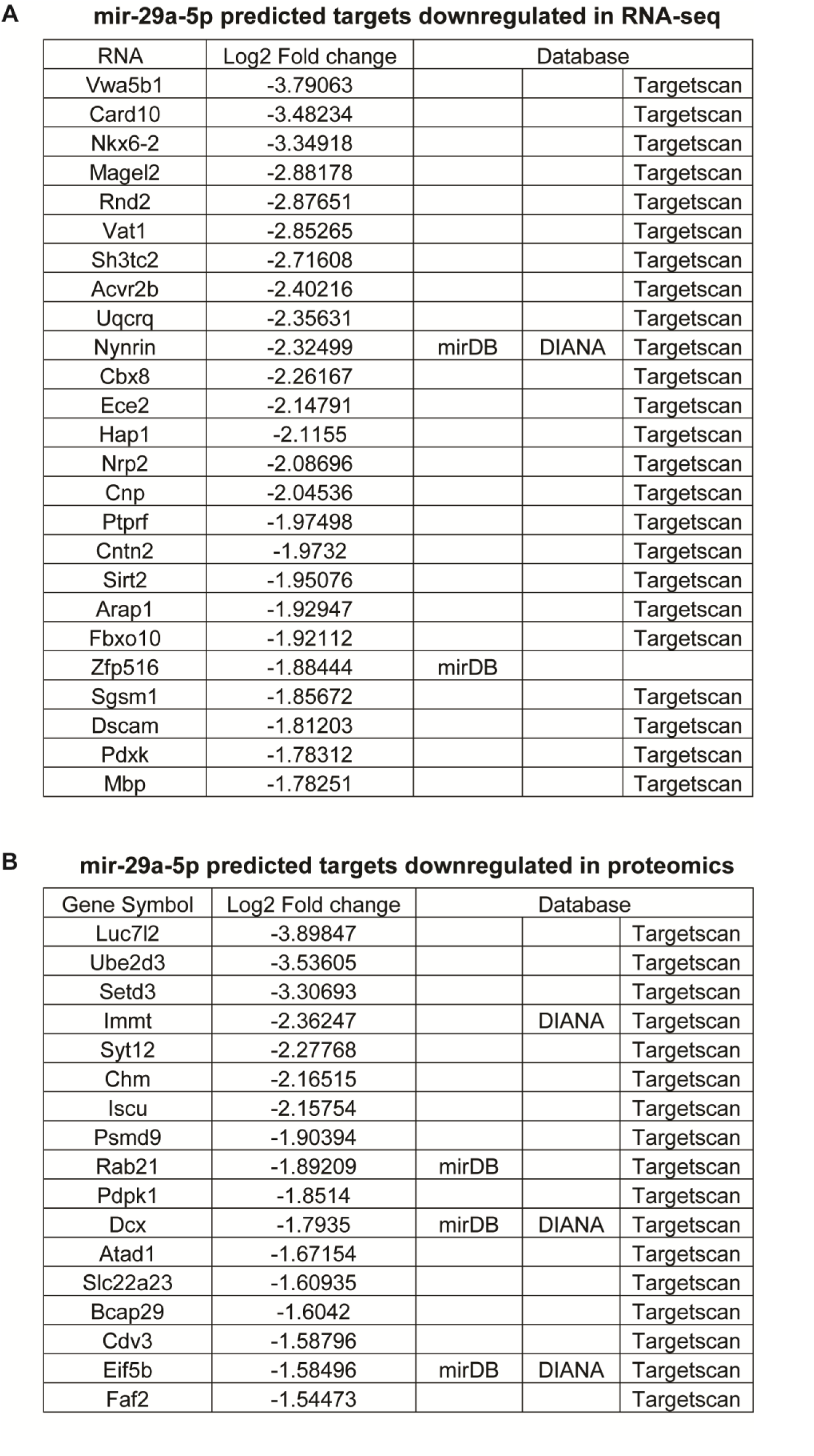
mir-29a-5p predicted targets are regulated in the transcriptome and proteome after acute restraint stress. Tables of the predicted mir-29a-5p targets that are downregulated at the RNA level (A) and protein level (B) in the BLC 30 days after an acute restraint session, based on RNA-seq and quantitative mass spectrometry. The database that identified each target is listed to the right of the log2 fold change values.

Ube2d3, a ubiquitin conjugating enzyme that was validated at both the protein (Fig 2E) and mRNA (Fig 3C) levels, is one of the most highly regulated targets of mir-29a-5p. The converging evidence of altered Ube2d3 levels in conjunction with mir-29a-5p regulation highlight the potential novel role of this miRNA pathway and specific target in the BLC’s stress-associated responses.

## Discussion

We have identified a molecular profile present in the BLC after a single acute stress paradigm and likely represents a long-term change in BLC signaling. The data indicate that acute restraint stress regulates the expression of miRNAs within the BLC, as well as mRNAs and proteins in both miRNA-dependent and -independent manners. The integration of multiple datasets at various stages of gene expression demonstrates low correspondence between mRNA and protein expression. This points to increased regulation at a step between transcription and translation and suggests that miRNAs may participate in the critical regulation of post-transcriptional processing to epigenetically modulate gene expression in response to a stressor. Although other studies have aligned miRNA expression with mRNA expression datasets in stress paradigms [32], our study represents the first that integrates such datasets with global proteomics from brain tissue to identify miRNA-mediated translational inhibition pathways affected long after a single stress event.

We have validated the increased expression of mir-29a-5p in the BLC 30 days after an acute restraint stress session, indicating that stress has a lasting effect on miRNA expression in a brain region essential for storage of long-term memories, stress responses and emotional regulation. Downstream of mir-29a-5p, we confirmed regulation of many mir-29a-5p putative targets at both the RNA and protein levels, highlighting the mir-29a-5p pathway as a key stress-responsive pathway in the BLC. In a more severe stress paradigm that lasts for 3 days, mir-29a was elevated 14 days later in the serum, but not the amygdala, of rodents stressed by restraint and foot shock [37]. mir-29c was decreased in serum immediately after the completion of the stress paradigm [37]. Interestingly, mir-29c was increased in the blood of human subjects three hours after a social stress paradigm, an elevation that positively correlated with an increased connectivity between the ventral medial prefrontal cortex and the insula [38]. Human mir-29c has high homology with mir-29a-5p and the two miRNAs are predicted to target many of the same genes [21; 39]. This suggests that stress many regulate the mir-29 family in both human and rodents in a conserved mechanism and that the stress-induced elevation is reflected beyond the brain, into the bloodstream. miRNAs can be released from neurons into serum viaextracellular vesicles, so it is reasonable to postulate that brain levels of mir-29a may influence those of the blood/serum and serve as a marker of stress responsiveness or stress history.

Other studies have reported miRNA changes in the hippocampus (mir-124a), central amygdala (mir-34) and basolateral amygdala after acute stress (mir-135a and mir-124) [26; 34; 40]. These studies collected tissue within 90 minutes of stress and identified immediate stress-responsive miRNAs that may have returned to baseline by the time tissue was collected in the current study, one month post-stress, for identification of long-lasting miRNA changes in the BLC. These changes may represent a neuroadaptation in response to the stressor, as they occurred 30 days after the stress event.

The RNA and protein datasets identified a number of significantly enriched pathways in the BLC that are associated with prior history of stress. While some of the signaling pathways such as MAPK (miRNA), calcium signaling (protein) and protein ubiquitination (mRNA) have been studied for their ability to activate cell signaling responses after a stress exposure, many other pathways were identified that contain molecules not typically studied in the stress field. Interestingly, the proteomics pathway analysis was more relevant to neurological processes than the RNA-seq pathway analysis. One of the validated upregulated genes was Gabrb2, which encodes Gamma-aminobutyric acid (GABA) A receptor, beta 2, a component of inhibitory GABA receptors. Given that increased inhibition of the BLC would be expected to dampen a stress response through reduced activation of the central amygdala’s motor output of a fear response, the long-term changes in Gabrb2 may represent a compensatory mechanism to mitigate responsivity to future stressors [10].

Ubiquitin conjugating enzyme e2 d3, Ube2d3, is a predicted target of mir-29a-5p and, consistent with translational suppression by miRNAs, was downregulated in stressed animals (mir-29a-5p was upregulated with stress). Since these protein changes were validated 30 days after the stress exposure, the downregulation of Ube2d3 likely represents a lasting neuroadaptation and lends supports to Ube2d3 being a genuine target of mir-29a-5p. Interestingly, mRNA levels of Ube2d3 were increased, which may be a homeostatic response to the protein downregulation of Ube2d3. Although there is little known about the role of this protein in the brain, it has been studied for its role in mediating oxidative stress and mitochondrial dynamics in neurodegenerative disorders [41; 42]. Since it regulates the rate of ubiquitination of proteins for proteosomal degradation, a downregulation of this protein may signify a decrease in the rate of protein turnover and be related to the expression differences observed in other proteins in the BLC after stress. In a chronic stress paradigm, stressed animals have reduced function of proteins essential for cognition and learning, the glutamate receptor subunits GluR1 and NR1, an effect that is mediated in part through stress-induced ubiquitin/proteosomal degradation of the subunits [43]. Ube2d3 coordinates with the E3 ubiquitin-protein ligases, one of which, Ube3a, has been implicated in plasticity and memory [44]. Additionally, the ubiquitin/proteasome system has been well studied in rodent models of fear learning for its role in memory consolidation, memory storage, plasticity and extinction learning [45]. Thus, Ube2d3 represents a new avenue to study downstream of mir-29a-5p signaling and highlights a pathway previously understudied in the stress field.

Studying the molecular consequences of acute stress is directly relevant to understanding mechanisms that confer increased vulnerability for development of psychiatric disorders. We have previously demonstrated that the same acute stress paradigm applied here in combination with auditory fear conditioning results in stress-enhanced fear learning, extinction resistance and development of PTSD-like behaviors and neuroadaptions [30]. Importantly, these effects were not present in animals that went through fear conditioning without prior restraint stress. Rather, an interaction between the acute stress experience and fear learning was required for the phenotype of enhanced stress-susceptibility. This work, in combination with the present study, further highlights the importance of understanding the processes by which acute stress experiences may alter brain function to put an individual at increased risk for adverse behaviors. Finally, since the lists of overlapping differentially expressed miRNAs with regulated proteins or RNAs do not include every RNA or protein differentially expressed, it is likely that a number of the expression changes observed may occur through miRNA-independent pathways. The data obtained here may serve as a resource for future studies into the molecular neuroadaptations that arise from a single stress event and have generated many novel hypotheses to examine in future studies.

## Materials and methods

### Animals

Adult male C57BL/6 mice, 8 weeks of age (The Jackson Laboratory, Bar Harbor, ME), were maintained on a 12:12 hour light/dark cycle and supplied with food and water ad libitum. Mice were group housed, acclimated to the facility for 1 week then handled for 3 days prior to experiments. All procedures associated with this study were approved by and performed in accordance with the guidelines of the Institutional Animal Care and Use Committee (IACUC) at The Scripps Research Institute, Florida and with national regulations and policies.

### Restraint stress

Acute restraint stress was performed using clear 50ml conical vials (Falcon Centrifuge Tubes) that had ten 5mm holes for ventilation. Animals were placed individually into their own tubes and restrained for 2 hours in a biosafety cabinet with overhead lights. Each tube was placed flat in an open box (120mm X 80mm X 50mm) to prevent minimize movement of the tube. Animals could hear and smell one another but not see each other during the procedure. Mice in nonstressed groups were briefly handled in another room during the restraint stress procedure. Following the procedure, all animals were returned to their homecage where they remained for 30 days. All animals were handled each week briefly for cage changes.

### RNA and protein extraction

After 60 minutes, 24 hours, or 30 days, animals were removed from their homecage, immediately anesthetized with isoflurane then rapidly decapitated. Brains were frozen in ice cold isopentane on dry ice then stored at -80°C until microdissection. For removal of the BLC, brains were dissected on dry ice in a brain block to maintain RNA integrity. For sequencing, qPCR and Western Blots, total RNA and proteins were simultaneously extracted from fresh frozen bilateral tissue punches using the miRVANA PARIS RNA extraction kit (Life Technologies, Carlsbad, CA) following the manufacturer’s instructions, as previously reported [46]. After the initial lysis step, tissue homogenates were split into RNA or protein tubes and processed separately. 5X radioimmunoprecipitation assay buffer (RIPA, 100mM Tris-HCl [pH 7.5], 750mM NaCl, 5mM ethylenediaminetetraacetic acid [EDTA], 5mM ethylene glycol-bis[β-aminoethyl ether]-N,N,N’,N’-tetraacetic acid, 5% nonyl phenoxypolyethoxylethanol, 5% sodium deoxycholate) with 5X Halt Protease and Phosphatase Inhibitor Cocktail (Thermo Scientific) was added to the protein fractions, which were sonicated, spun at 4°C for 10 min at 15,000 X g to remove cellular debris and then analyzed for yield with the Thermo Scientific bicinchoninic acid (BCA) Assay (Thermo Scientific). RNA concentrations were measured with a Qubit 3.0 Fluorometer and the Qubit RNA High Sensitive Assay (Invitrogen, Carlsbad, California, USA).

### mRNA Library Preparation, Sequencing and data analysis

Preparation of mRNA libraries, mRNA sequencing and analysis of mRNA expression data was performed as previously described [30]. Individual samples of BLC tissue were sequenced 30 days after stress (naïve N=2, stressed N=4) and significantly regulated mRNAs were identified that had an uncorrected p-value of less than 0.05.

### miRNA Library Preparation, Sequencing and data analysis

Small-RNA sequencing was performed on two biological replicate cohorts of samples at two distinct sequencing facilities, the Scripps Florida Genomics Core (N=3/group) and the Genomic Sequencing Laboratory at Hudson Alpha (N=4/group). For sequencing at Scripps, total RNA was quantified by Qubit and run on the Agilent 2100 Bioanalyzer (Agilent Technologies, Santa Clara, CA) for quality assessment. All RNA Integrity Numbers (RINs) were > 7.0. 750ng total RNA was processed using the TruSeq small RNA library prep kit (Illumina, San Diego, CA). Briefly, RNA samples were sequentially ligated with 3’ and 5’ RNA adapters. The adapter ligated RNA molecules were reverse transcribed and PCR amplified. The polymerase chain reaction (PCR) amplified products were visualized on a bioanalyzer and pooled 1:1 based on the concentration of the miRNA library peaks (migrating between 130-170bp). The pooled libraries were size selected on a 6% tris-borate EDTA gel to recover the miRNA library fragments. After gel extraction and ethanol precipitation of the DNA, the final library was quantified using Qubit and visualized on the Bioanalyzer. The library pool was loaded onto the NextSeq 500 flow cell (Illumina) at 1.8pM final concentration and sequenced with single-end 50bp sequencing to yield at least 20 million reads per sample. Reads were trimmed of adapter sequences using cutadapt [47] with the following options: -a TGGAATTCTCGGGTGCCAAGG -O 3 -m 14 -M 25. All known hairpin and mature miRNAs were downloaded from miRbase version 21 [20; 21; 39; 48]. miRNAs were identified using mirDeep2 [19] against the mouse genome (GENCODE version GRCm38.p3) using mapper.pl with the following options: mapper.pl cutadapt.fastq -e -h -p mouseGENCODE -s mapped.fa -t samp.arf -m –v, followed by quantifier.pl using the options: -p precursor.fa -m mature.fa -r mapped.fa -t mmu, and finally by the miRDee2.pl wrapper script with the options: mapped.fa Genome.fasta samp.arf MaturemiRNA.fa none HairpinPrecursor.fa-t Mouse -q miRBase.mrd. Tables of counts were generated using unique combinations of the miRNA precursor and mature miRNA names and analyzed using DESeq2 (version 1.10.1, R version 3.2.3) [49]. For sequencing performed at Hudson Alpha, 200ng total RNA from each sample was taken into a small RNA library preparation protocol using NEBNext Small RNA Library Prep Set for Illumina (New England BioLabs Inc., Ipswich, MA, USA) according to the manufacturer’s protocol. Briefly, 3` adapters were ligated to total input RNA followed by hybridization of multiplex SR reverse transcription (RT) primers and ligation of multiplex 5` SR adapters. RT was done using ProtoScript II RT for 1 hour at 50°C. Immediately after RT reaction, PCR amplification was performed for 15 cycles using LongAmp Taq 2X master mix. Illumina indexed primers were added to uniquely barcode each sample. Post-PCR material was purified using QIAquick PCR purification kit (Qiagen Inc., Valencia, CA, USA). Post-PCR yield and concentration of the prepared libraries were assessed using Qubit and DNA 1000 chip on Agilent 2100 Bioanalyzer (Applied Biosystems, Carlsbad, CA, USA), respectively. Size selection of small RNA was done using a 3% dye free agarose gel cassette on Pippin prep instrument (Sage Science Inc., Beverly, MA, USA). Post-size selection yield and concentration of the libraries were assessed using Qubit 2.0 Fluorometer and DNA High sensitivity chip on Agilent 2100 Bioanalyzer, respectively. Accurate quantification for sequencing applications was performed using the qPCR-based KAPA Biosystems Library Quantification kit (Kapa Biosystems, Inc., Woburn, MA, USA). Each library was diluted to a final concentration of 1.25nM and pooled in equimolar ratios prior to clustering. Single End (SE) sequencing (50bp) was performed to generate at least 15 million reads per sample on an Illumina HiSeq2500 sequencer (Illumina). Post processing of the sequencing reads from miRNA-seq experiments from each sample was performed as per the Genomic Services Laboratory’s unique in-house pipeline. Briefly, quality control checks on raw sequence data from each sample were performed using FastQC (Babraham Bioinformatics, London, UK). Raw reads were imported on a commercial data analysis platform AvadisNGS (Strand Scientifics, CA). Adapter trimming was done to remove ligated adapter from 3’ end of the sequenced reads with only one mismatch allowed, poorly aligned 3’ ends were also trimmed. Sequences shorter than 15 nucleotides length were excluded from further analysis. Trimmed Reads with low qualities (base quality score less than 30, alignment score less than 95, mapping quality less than 40) were removed. Filtered reads were then used to extract and count the small RNA which was annotated with miRNAs from the miRBase release 20 database [20; 21; 39; 48]. The quantification operation carries out measurement at both the gene level and at the active region level. Active region quantification considers only reads whose 5’ end matches the 5’ end of the mature miRNA annotation. miRNAs that had at least 0.5 log2 fold change between SS and SR in both sequencing runs were considered ‘candidate miRNAs’ and analyzed for pathway analysis with DIANA mirPATH software [50], which integrates information from TargetScan [17] to predict significant pathways of enriched target genes for a give list of miRNAs. Target prediction analysis was examined using DIANA’s microCts and Tarbase algorithms as well as TargetScan.

### Immunoblotting

Five micrograms of purified protein was analyzed on Bio-Rad 4-20% gels as previously described[51]. A primary antibody directed against Ube2d3 (1:5000 dilution; catalog# Ab176598, Abcam, Cambridge, MA) was incubated overnight at 4°C. Horseradish peroxidase conjugated secondary antibodies were (1:2000 dilution; Promega, Madison, WI) incubated with blots for two hours at room temperature. Total protein stains (Pierce Reversible Protein Stain Kit, Thermo Scientific) were obtained after each transfer and used as a loading control for normalization in Image J Software[52].

### QPCR validation

For validation of smRNA-seq, a cDNA library was created from 50ng of total RNA using the mirCURY LNA RT Kit (Qiagen, Germantown, MD) following the manufacturer’s protocol and technical replicate samples that were sequenced. cDNA was diluted 1:80 for PCR. PCR reactions were performed in triplicate for each sample using the miRCURY LNA SYBR Green PCR Kit and the following locked nucleic acid (LNA) SYBR green primers from Qiagen (formerly Exiqon): mir-135b-3p, YP00204427; mir-199a-5p, YP00204494; mir-29a-5p, YP00204430. PCR cycle conditions were 95°C for ten minutes then 40 amplification cycles of 95°C for 10 seconds to denature and 60°C for 1 minute to anneal and elongate. A melt curve was run for each primer to verify the formation of only one product. For validation of RNA-seq, cDNA synthesis and qPCR were performed as previously described[46] with 50ng of RNA using the following Taqman Assays (Thermo-Life Scientific): Chm, Mm00517015_m1; Dnajb4, Mm00508908_m1; Gabrb2, Mm00433467_m1; Rpl15, Mm01727317_g1; Slc8a1, Mm01232254_m1; Ube2d3, Mm00787086_s1. Data were normalized to the housekeeping snoRNA snord68 using the ΔΔc_t_ method [53].

### Quantitative mass spectrometry

30 days after the stress procedure, animals were removed from their homecage, immediately anesthetized with isoflurane then rapidly decapitated. Brains were frozen in ice cold isopentane on dry ice then stored at -80°C until microdissection. For removal of the BLC, brains were dissected on dry ice in a brain block to maintain RNA integrity. Frozen BLC tissue punches were pooled (8 animals/group). Pooled samples from each treatment group were submitted for liquid chromatography–mass spectrometry (LC-MS) that was performed on an LTQ Orbitrap Elite (Thermo Fisher, Waltham, MA) equipped with Waters NanoAcquity HPLC pump (Milford, MA) at the Harvard Mass Spectrometry and Proteomics Resource Laboratory. Peptides were separated onto a 100 μm inner diameter microcapillary trapping column packed first with approximately 5 cm of C18 Reprosil resin (5 μm, 100 Å, Dr. Maisch GmbH, Germany) followed by analytical column ∼20 cm of Reprosil resin (1.8 μm, 200 Å, Dr. Maisch GmbH, Germany). Separation was achieved through applying a gradient from 5–27% ACN in 0.1% formic acid over 90 min at 200 nl min-1. Electrospray ionization was enabled through applying a voltage of 1.8 kV using a home-made electrode junction at the end of the microcapillary column and sprayed from fused silica pico tips (New Objective, MA). The LTQ Orbitrap Elite was operated in data-dependent mode for the mass spectrometry methods. The mass spectrometry survey scan was performed in the Orbitrap in the range of 395 –1,800 m/z at a resolution of 6 × 104, followed by the selection of the twenty most intense ions (TOP20) for CID-MS2 fragmentation in the Ion trap using a precursor isolation width window of 2 m/z, AGC setting of 10,000, and a maximum ion accumulation of 200 ms. Singly charged ion species were not subjected to CID fragmentation. Normalized collision energy was set to 35 V and an activation time of 10 ms. Ions in a 10 ppm m/z window around ions selected for MS2 were excluded from further selection for fragmentation for 60 s. The same TOP20 ions were subjected to HCD MS2 event in Orbitrap part of the instrument. The fragment ion isolation width was set to 0.7 m/z, AGC was set to 50,000, the maximum ion time was 200 ms, normalized collision energy was set to 27V and an activation time of 1 ms for each HCD MS2 scan. Raw data were submitted for analysis in Proteome Discoverer 2.1.0.81 (Thermo Fisher) software. Assignment of MS/MS spectra was performed using the Sequest HT algorithm by searching the data against a protein sequence database including all entries from the Human Uniprot database (SwissProt 16,768 and TrEMBL 62,460 total of 79,228 protein forms, 2015) [54] and other known contaminants such as human keratins and common lab contaminants. Sequest HT searches were performed using a 20 ppm precursor ion tolerance and requiring each peptide’s N-/C termini to adhere with Trypsin protease specificity, while allowing up to two missed cleavages. 6-plex tandem mass tags (TMT) on peptide N termini and lysine residues (+229.162932 Da) was set as static modifications while methionine oxidation (+15.99492 Da) was set as variable modification. A MS2 spectra assignment false discovery rate (FDR) of 1% on protein level was achieved by applying the target-decoy database search. Filtering was performed using a Percolator (64bit version [55]). For quantification, a 0.02 m/z window centered on the theoretical m/z value of each the six reporter ions and the intensity of the signal closest to the theoretical m/z value was recorded. Reporter ion intensities were exported in result file of Proteome Discoverer 2.1 search engine as an excel tables. The total signal intensity across all peptides quantified was summed for each TMT channel, and all intensity values were adjusted to account for potentially uneven TMT labeling and/or sample handling variance for each labeled channel. Candidate proteins were identified as those that had at least 1.5 log2fold change between treatment groups.

## Acknowledgments

We thank the Scripps Florida Genomics Core for sequencing services, Nripesh Prasad at the Genomic Services Lab at Hudson Alpha for sequencing services and data analysis, and Adrian Reich and the Bioinformatics Core for data analysis.

## Supporting information

Table SE.1

Table SE.2

Table SE.3

Table SE.4

